# Efficient biallelic knock-in in mouse embryonic stem cells by *in vivo*-linearization of donor and transient inhibition of DNA Polymerase θ/DNA-PK

**DOI:** 10.1101/2021.05.30.446338

**Authors:** Daisuke Arai, Yoichi Nakao

**Affiliations:** School of Advanced Science and Engineering, Waseda University, 3-4-1 Okubo, Shinjuku-ku, Tokyo 169-8555, Japan; Research Institute for Science and Engineering, Waseda University, 3-4-1 Okubo, Shinjuku-ku, Tokyo 169-8555, Japan

## Abstract

CRISPR/Cas9-mediated homology-directed repair (HDR) is used for error-free targeted knock-in of foreign donor DNA. However, the low efficiency of HDR-mediated knock-in hinders establishment of knock-in clones. Double-strand breaks (DSBs) induced by CRISPR/Cas9 are preferentially repaired by non-homologous end joining (NHEJ) or microhomology-mediated end joining (MMEJ) before HDR can occur, thereby preventing HDR-mediated knock-in. NHEJ/MMEJ also cause random integrations, which give rise to false-positive knock-in events, or silently disrupt the genome. In this study, we optimized an HDR-mediated knock-in method for mouse embryonic stem cells (mESCs). We succeeded in improving efficiency of HDR-mediated knock-in of a plasmid donor while almost completely suppressing NHEJ/MMEJ-based integration by combining *in vivo*-linearization of the donor plasmid, transient knockdown of DNA Polymerase θ, and chemical inhibition of DNA-dependent protein kinase (DNA-PK) by M3814. This method also dramatically improved the efficiency of biallelic knock-in; at the *Rosa26a* locus, 95% of HDR-mediated knock-in clones were biallelic. We designate this method BiPoD (<u>Bi</u>allelic knock-in assisted by <u>Po</u>l θ and <u>D</u>NA-PK inhibition). BiPoD achieved simultaneous efficient biallelic knock-in into two loci. BiPoD, therefore, enables rapid and easy establishment of biallelic knock-in mESC lines.

## Introduction

Genomic integration of foreign donor DNA through homology-directed repair (HDR), exploited for classical gene-targeting technology, has been dramatically facilitated by development of the CRISPR/Cas9 system [1]. In this system, Cas9 endonuclease and single guide RNA (sgRNA) induce double-strand breaks (DSBs) at specific target sites into which donor DNA can be knocked-in. The broken sites are repaired by HDR using foreign donor DNA as a template, resulting in error-free targeted integration of the donor. HDR-mediated integration enables precise knock-in to be performed, including donor-based gene knock-out, tagging of endogenous genes, visualization of genomic regions, and insertion of expression cassettes into defined loci such as safe-harbor regions [2].

Foreign donor DNA can be integrated into a host genome using any of several host DSB repair mechanisms. Therefore, to efficiently obtain HDR-mediated knock-in cells it is essential to reduce other types of integration events. Non-homologous end joining (NHEJ) is a major pathway for HDR-independent genomic integration, in which the ends of linearized donor DNA are directly ligated to the broken ends of genomic DNA [3]. This process is executed by multiple machineries, including DNA-dependent protein kinase (DNA-PK) and DNA ligase IV (LIG4) [4, 5]. Another pathway is microhomology-mediated end joining (MMEJ). In contrast to NHEJ, the ends of donor and genomic DNA are aligned via microhomologous regions (≥ 1 bp) before joining [6]. Recent studies have shown that DNA polymerase theta (Pol θ or POLQ) plays a central role in MMEJ, thus this pathway is also called polymerase theta-mediated end joining [7, 8]. Donor DNA is efficiently integrated into CRISPR/Cas9-induced DSB sites (on-target) through NHEJ or MMEJ with or without indel mutation [9, 10]. Donor DNA can also be integrated into non-targeted sites (off-target) by way of NHEJ or MMEJ; this is called random integration [11–13]. Random integration is the easiest way to obtain transgenic cells, but the integration is unregulated resulting in unwanted genomic disruption and the generation of false-positive cells that incorporate marker genes at incorrect sites.

An advantage of HDR-mediated integration over NHEJ/MMEJ-based methods is the availability of negative selection markers, which are located outside the donor DNA homology arms and are only expressed if the entire donor is integrated through NHEJ/MMEJ. Frequently, however, only part of the donor is integrated into on- or off-target sites of the genome. In addition to donor DNA, part of the Cas9/sgRNA expression vector can also be integrated [14]. If negative selection markers are not integrated, it is very hard to detect the occurrence of such integrations. These “silent integrations” disrupt the genome but are difficult to avoid through donor DNA design features. Because silent integrations occur by NHEJ and/or MMEJ, suppressing NHEJ/MMEJ events is of great importance to obtain scarless knock-in cells.

HDR is a rare event even when the CRISPR/Cas9 system is employed; therefore, it is difficult to obtain HDR-mediated knock-in cells [15, 16]. The success rate for biallelic knock-in is extremely low, although it is required for various experimental strategies. Classically, two donor DNA templates with different marker genes [e.g., puromycin/blasticidin (BS) resistance genes and EGFP/mCherry] have been used to select biallelic knock-in cells [17, 18], but such dual markers are not always available. Furthermore, random integration of the antibiotic resistant genes can give rise to false-positive cells and prevent the acquisition of cells with precise biallelic knock-in. Thus, it would be advantageous to improve knock-in efficiency. In addition to HDR using foreign donor DNA as a template, DSBs induced by CRISPR/Cas9 are repaired by various other pathways, as described above. Once a site is repaired with accompanying genomic changes, such as indel mutations or donor insertions, the site is no longer a target of the same sgRNA/Cas9 complex. Therefore, HDR-mediated knock-in must occur before the target site is repaired by other mechanisms. Manipulating NHEJ and MMEJ pathways by chemical inhibitors, as well as overexpression and/or knockdown of pathway genes, have been tested in a large number of studies to improve HDR-mediated knock-in efficiency [19–21]. However, results from different studies have been inconsistent and a gold-standard method has not yet been established. Accordingly, biallelic knock-in of long donor DNA remains difficult and time-consuming.

Here, we show an optimized method for HDR-mediated knock-in applicable to mouse embryonic stem cells (mESCs). Combining *in vivo*-linearization of donor DNA, knockdown of the *Polq* gene, and chemical inhibition of DNA-PK, we achieved specific HDR-mediated knock-in with very high efficiency of biallelic integration. Our method, termed BiPoD (<u>Bi</u>allelic knock-in assisted by <u>Po</u>l θ and <u>D</u>NA-PK inhibition), provides an easy and rapid way to obtain precise knock-in mESCs.

## Results

### *In vivo*-linearization of a plasmid donor improves the efficiency of HDR-mediated knock-in

To evaluate the efficiency of HDR-mediated knock-in, *Rosa26a*, a widely used safe harbor locus in the mouse genome, was adopted as a target. The donor is shown in Fig. 1a. The CAG promoter-driven *mCherry* gene was inserted between 5′ and 3′ homology arms (0.8 kb and 0.9 kb, respectively), while an *EGFP* expression cassette was located downstream of the 3′ arm. If the donor is precisely integrated through HDR, the cells exhibit only red fluorescence (mCherry-positive/EGFP-negative, R+G−). The cells exhibit both red and green fluorescence if the entire donor sequence is integrated through a mechanism other than HDR, irrespective of integration site (mCherry-positive/EGFP-positive, R+G+). Another possibility is "partial integration", in which only part of the donor sequence is integrated into either *Rosa26a* or random loci. The fluorescence pattern depends on the region of the donor that is integrated. Cells are R+G+ if a majority of the donor is integrated, including both *mCherry* and *EGFP* expression cassettes, while integration of donor sequence fragments leads to R+G−, R−G+, or even R−G− cells, depending on which fragments are integrated. Here, we have to pay attention to partial integration. R+G− derived from partial integration could not be distinguished from that derived from legitimate HDR. Additionally, integration could not be detected if neither the *mCherry* nor *EGFP* expression cassette was included. We hypothesized that the occurrence of partial integration can be roughly estimated from the rate of R−G+ (only the *EGFP* expression cassette is integrated) because all partial integration events are thought to be caused by common mechanisms.

**Figure 1.**
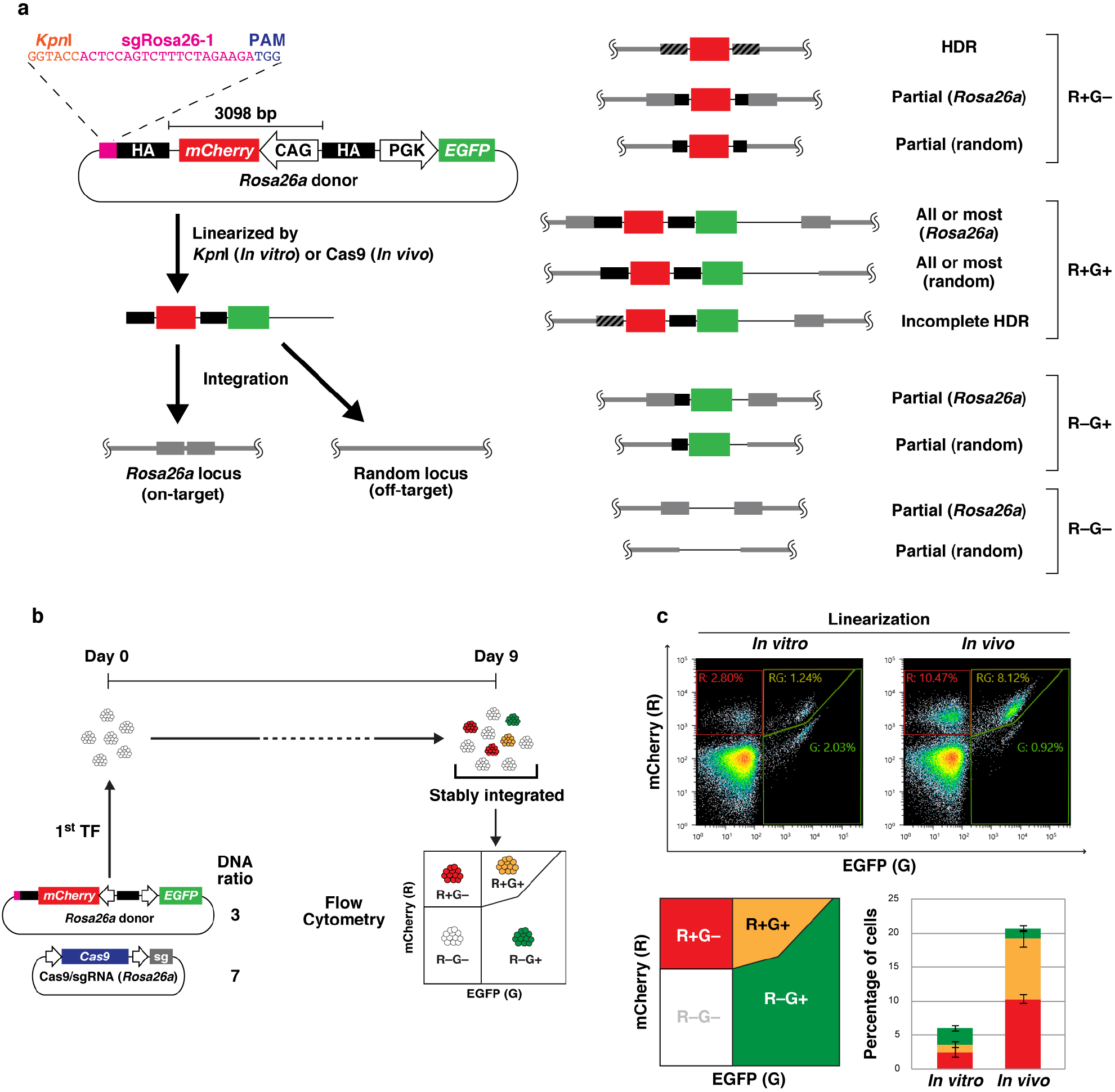
Comparison of methods to linearize plasmid donors. (a) Schematic illustration of the *Rosa26a* donor and possible modes of integration. Wide black bars and wide gray bars indicate the homology arms in the donor and the homologous region in the genome, respectively. Black-and-white striped bars denote the homologous region undergoing HDR. The pink bar indicates the sgRNA target sequence (sgRosa26-1). (b) Experimental scheme. Weight ratio of donor DNA and Cas9/sgRNA expression vector is shown. (c) Flow cytometry analysis of mESCs transfected with the *Rosa26a* donor and Cas9/sgRNA expression vector. A series of experiments, from transfection to flow cytometric analysis, was independently performed in triplicate. Scatter plots show representative results from three experiments. The average ± standard deviation percentages of R+G− (red), R+G+ (yellow), and R−G+ (green) cells are shown in the stacked bar graph.

First, we compared the knock-in efficiency of *in vivo*- and *in vitro*-linearized plasmid donor DNA. To this end, a *Kpn*I site and sgRosa26-1 sequence (a previously validated CRISPR/Cas9 target [22]) were located upstream of the 5′ arm (Fig. 1a). mESCs were transfected with the same amount of circular or *Kpn*I-digested donor DNA. The circular donor is digested by Cas9 in the transfectants (*in vivo*-linearized), while the donor pre-digested by *Kpn*I (*in vitro*-linearized) is directly used as an HDR template or integrated through other mechanisms. After simultaneous transfection of the donors with the expression vector of Cas9 and the sgRNA against *Rosa26a*, mESCs were cultured for 9 days to completely screen out transient expression of mCherry and/or EGFP and were then analyzed by flow cytometry (Fig. 1b, c). Only 2.5% of the *in vitro*-linearized donor transfectants became R+G−. However, the percentage of R−G+ transfectants was 2.4%, indicating that some or most of the R+G− cells occurred by partial integration and that the true occurrence of HDR-mediated knock-in was much lower than 2.5%. The percentage of R+G− cells was much higher when the *in vivo*-linearized donor was used (10%, *p* = 1.5 × 10^−4^vs *in vitro*-linearization), while that of R−G+ cells was only 1.4%. These results indicated that the *in vivo*-linearization of the donor significantly increased the occurrence of HDR-mediated knock-in, although partial integration still occurred at a non-negligible level. In addition, the percentage of R+G+ cells resulting from non-HDR integration, was also increased when using the *in vivo*-linearized donor (1.1% to 8.9%, *p* = 5.0 × 10^−4^).

### Combination of transient *Polq* knockdown with inhibition of DNA-PK decreases undesired integration of donors

To take advantage of the efficient HDR resulting from *in vivo*-linearization, we attempted to suppress non-HDR integration events during knock-in using *in vivo*-linearized donor. Random integration never occurs in *POLQ* and *LIG4* double knock-out cells [12, 13]. Pharmacological inhibition of DNA-PK by NU7441 in *Polq* KO mESCs increased the ratio of HDR to random integration to 1:1 at best [13]. Irreversible knock-out is not suitable for general knock-in experiments; therefore, we examined whether transient inhibition of Pol θ and DNA-PK in combination with *in vivo*-linearization of donor suppressed non-HDR integration events.

A number of small compounds that specifically and potently inhibit DNA-PK are available, whereas few specific Pol θ inhibitors have been reported. We, therefore, transiently knocked down Pol θ using shRNA. A series of experiments was performed as shown in Fig. 2a. To take into account the time required for repression by shRNA, mESCs were transfected with plasmids expressing shRNA against *Polq* or scrambled shRNA and a puromycin-resistance gene one day before they were transfected with the circular plasmid donor targeting *Rosa26a* and the Cas9/sgRNA expression vector. These cells were treated with puromycin to select *Polq*-knockdown cells. (Fig. 2b). The cells were also treated with a DNA-PK inhibitor for 3 days after transfection of the donor and Cas9/sgRNA vectors. After culture for 6 days (10 days from 1st transfection), the cells were analyzed by flow cytometry.

**Figure 2.**
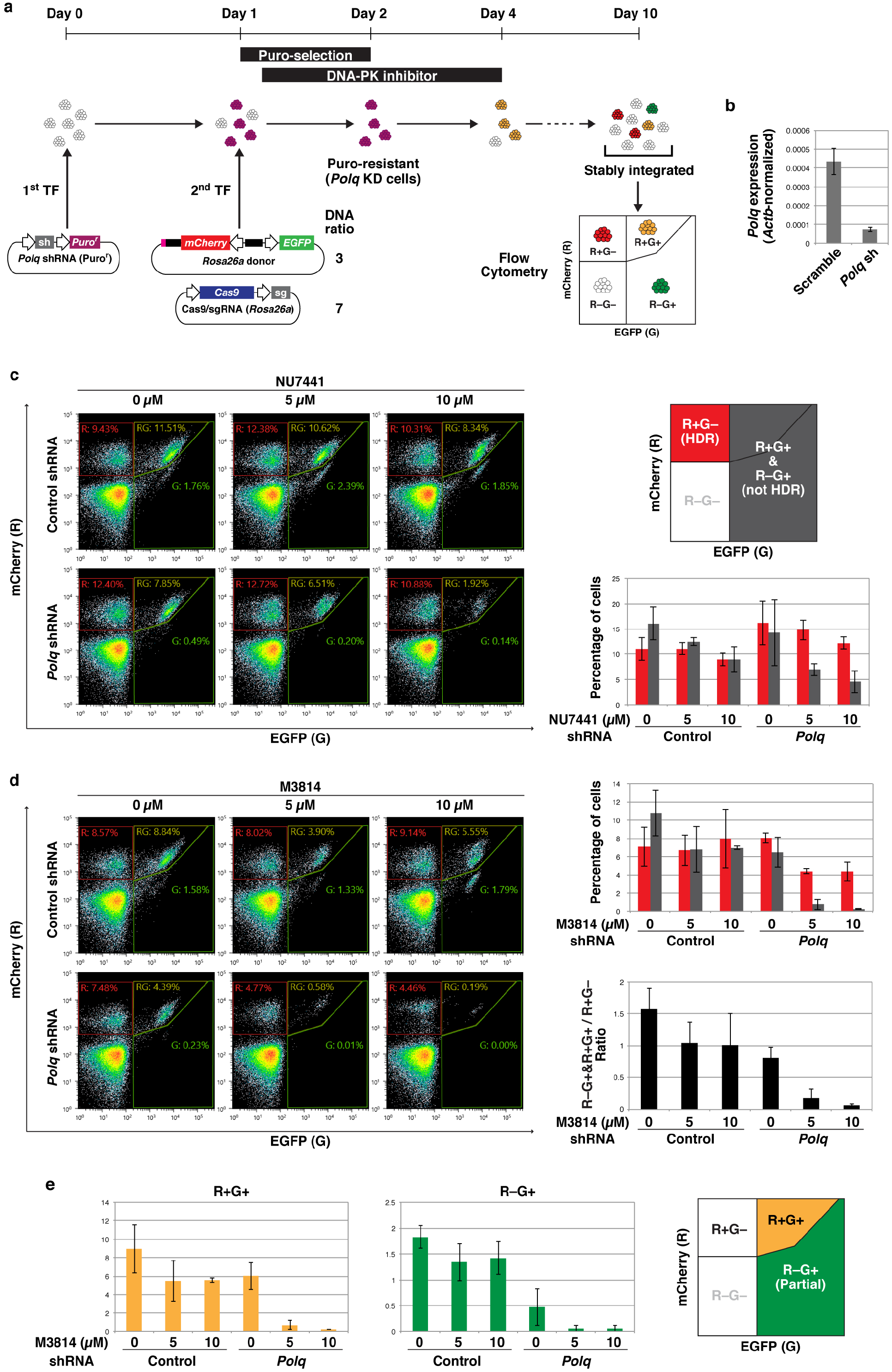
Effects of transient *Polq* knockdown combined with DNA-PK inhibitors on HDR-mediated knock-in. (a) Experimental scheme. Weight ratio of donor DNA and Cas9/sgRNA expression vector is shown. (b) Expression levels of *Polq* in mESCs 72 hours after transfection of *Polq* or scrambled shRNA were determined by qRT-PCR. Successfully transfected cells were selected by puromycin treatment. The bars represent the average ± standard deviation of triplicate qPCR samples normalized against *Actb*. (c, d) Flow cytometry analysis of mESCs transfected with the shRNA, *Rosa26a* donor, and Cas9/sgRNA expression vector. A series of experiments, from transfection to flow cytometric analysis, was independently performed in triplicate. Scatter plots show representative results from the three experiments. The average ± standard deviation percentages of R+G− (red) and R+G+ & R−G+ (gray) cells are shown in the bar graph. A black bar graph represents the ratio of R+G+ & R− G+ vs R+G−. (e) Percentage of R+G+ (yellow) and R−G+ (green) cells in the experiments shown in d. The average ± standard deviation is shown in the bar graphs.

Initially, we examined the effect of NU7441 in combination with *in vivo*-linearization of donor DNA and transient knockdown of *Polq* (Fig. 2c). NU7441 was added to mESCs at 0 (DMSO only), 5, or 10 μM, because NU7441 is moderately toxic and no mESCs survived with 20 μM NU7441. Knockdown of *Polq* alone did not significantly change the occurrence of R+G− (mainly HDR) or R+G+ & R−G+ (non-HDR) cells. *Polq* knockdown + 10 μM NU7441 decreased the proportion of R+G+ & R−G+ cells from 16% to 4.5% (3.5-fold, *p* = 0.011), and the R+G+ & R−G+/R+G− ratio from 1.47 to 0.37 (*p* = 7.2 × 10^−5^). These results show that simultaneous transient inhibition of Pol θ by shRNA and DNA-PK by NU7441 is able to suppress non-HDR donor integration to some extent.

M3814 is a novel DNA-PK inhibitor, which can enhance HDR efficiency [23, 24]. M3814 inhibits NHEJ more strongly than NU7441 in human induced pluripotent stem cells [25] but there are no reports of using M3814 in combination with Pol θ inhibition. We examined the efficacy of M3148 instead of NU7441 in our scheme (Fig. 2d). We used M3814 at up to 10 μM because, similarly to NU7441, it exhibited moderate toxicity to mESCs. Transient treatment with 10 μM M3814 decreased the number of living cells, particularly in combination with *Polq* knockdown, but the surviving cells grew and formed dome-shaped colonies, a characteristic of undifferentiated mESCs (Fig. S1). Treatment with M3814 alone did not significantly affect the percentage of R+G+ & R− G+ cells (non-HDR integration). However, a combination of M3814 and *Polq* knockdown dramatically reduced the percentage of R+G+ & R−G+ cells. In particular, *Polq* knockdown + 10 μM M3814 decreased the percentage of R+G+ & R−G+ cells from 11% to 0.25% (43-fold, *p* = 4.5 × 10^−5^). Although the percentage of R+G− cells tended to drop non-significantly (7.1% to 4.4%), the R+G+ & R−G+/R+G− ratio was strikingly lower than that of control cells (1.57 vs 0.06, *p* = 3.9 × 10^−4^). These results indicate that in knock-in experiments using *in vivo*-linearized donor, and a combination of M3814 and *Polq* shRNA almost eliminates non-HDR integration events, without preventing HDR.

Knockdown of *Polq* alone markedly reduce the occurrence of R−G+ cells (1.8% to 0.47%, 3.9-fold, *p* = 3.8 × 10^−4^), although it did not significantly affect the occurrence of R+G+ cells (8.9% to 6.0%, 1.5-fold) (Fig. 2e). M3814 alone had no significant effect on the occurrence of R−G+ cells, even at 10 μM (Fig. 2e). Therefore, inhibition of Pol θ is essential for the decreased numbers of R−G+ cells, i.e., partial integration of donor DNA, and M3814 enhances the effect of Pol θ inhibition. These results indicated that the slight decrease in the percentage of R+G− cells by *Polq* knockdown with 5 or 10 μM M3814 (Fig. 2d) resulted from fewer false-positives caused by partial integrations.

### Combination of *Polq* knockdown with M3814 treatment leads to biallelic knock-in

We focused on the fluorescence intensity of mCherry in the R+G− population (Fig. 3a and Fig. S2). The control cells (scrambled shRNA + DMSO) showed a single fluorescence peak, while a second peak emerged in some conditions. There was an approximately two-fold difference between the intensity of these two peaks, indicating that the weaker peak and the stronger peak corresponded to cells harboring one and two copies of the *mCherry* gene, respectively. Given that the *mCherry* genes in the R+G− cells were mainly integrated through HDR, these two peaks were thought to be derived from monoallelic and biallelic integration of the *mCherry* gene into *Rosa26a*. The stronger peak was significantly increased by *Polq* shRNA + 10 μM NU7441, while the weaker peak was almost completely abolished by *Polq* shRNA + 10 μM M3814. These results indicate that simultaneous inhibition of Pol θ and DNA-PK improves biallelic knock-in efficiency. To confirm this speculation, clones isolated from the R+G− populations were analyzed by genomic PCR (Fig. 3b). Only 6% (1/16) of control cell clones (scrambled shRNA + DMSO) harbored biallelic *mCherry* genes in *Rosa26a*. However, 33% (8/24) and 95% (20/21) of clones were biallelic for *Rosa26a mCherry* in *Polq* shRNA + 10 μM NU7441 and *Polq* shRNA + 10 μM M3814 cells, respectively. These results were in good agreement with the fluorescence intensity patterns observed by flow cytometry analysis (Fig. 3a). This method, using *in vivo*-linearized donor, *Polq* shRNA, and 10 μM M3814, was designated as BiPoD (<u>Bi</u>allelic knock-in assisted by <u>Po</u>l θ and <u>D</u>NA-PK inhibition).

**Figure 3.**
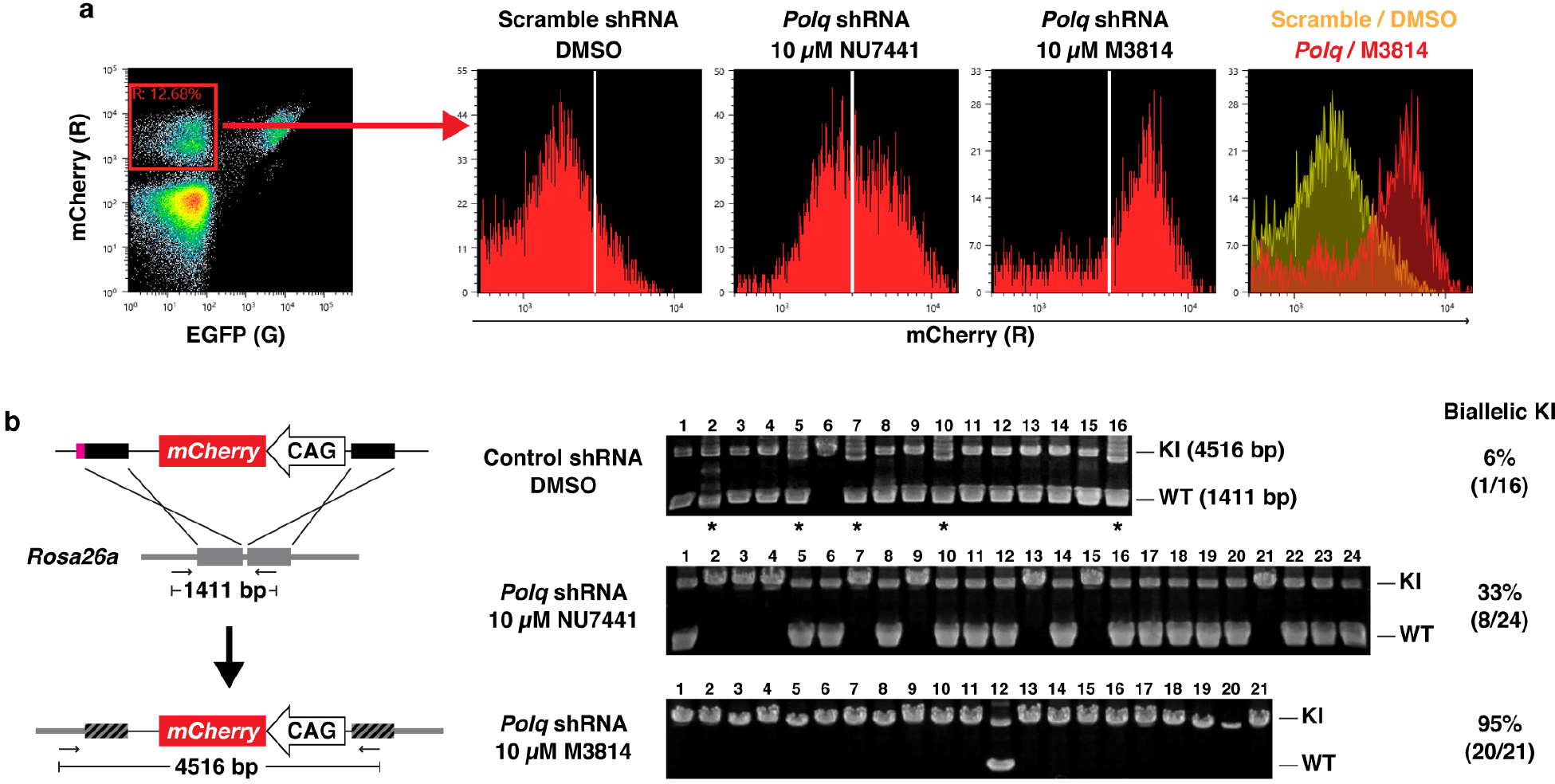
Biallelic knock-in of *mCherry* into *Rosa26a* promoted by *Polq* knockdown and DNA-PK inhibition. (a) Fluorescence intensities of mCherry in R+G− populations from the experiments shown in Fig. 2c and d. Representative histograms from three experiments are shown. (b) Genomic PCR analysis of clones isolated from R+G− populations. Left, position of primers and the size of amplicons are shown. Right, the PCR products were analyzed on agarose gels. Positions of knock-in allele (KI) and wild-type allele (WT) are indicated. Asterisks indicate clones in which PCR bands of unexpected size were observed.

### Combination of *Polq* knockdown with M3814 treatment can be used to tag endogenous genes

BiPoD was tested for its ability to tag the endogenous *Sox2* gene, which encodes a nuclear-localized transcription factor that plays important roles in mESCs. A donor was designed to insert an EGFP-mAID tag at the C-terminal end of *Sox2*. Similar to the *Rosa26a* donor, a CRISPR target sequence was located upstream of the 5′ homology arm for *in vivo*-linearization. The BS-resistance gene (*BS^r^*) under control of the PGK promoter was integrated downstream of the EGFP-mAID tag sequence, and an *mCherry* expression cassette was located outside the 3′ homology arm to detect non-HDR integration events (Fig. 4a). mESCs were transfected with this donor together with a Cas9/sgRNA expression vector, and the transfectants were cultured until transient expression was lost. In the EGFP-positive (R−G+) cells, EGFP was mostly localized in the nucleus, indicating that EGFP was expressed as a fusion protein with SOX2 (Fig. 4b). The EGFP-mAID tag is promoter-less; therefore, this donor is expected to express only mCherry (R+G−) when the whole donor sequence is non-HDR integrated, or in the case of partial integrations. A minor fraction of cells expressed both nuclear-localized EGFP and diffused mCherry (R+G+) (Fig. 4c). This likely resulted from incomplete HDR, simultaneous occurrence of both HDR- and non-HDR-mediated integrations, or fusion with other proteins because the homology arm included sequence encoding the nuclear localization signal of SOX2 [26]. In comparison to the control conditions, the percentage of R+G+ & R+G− cells (non-HDR) was drastically decreased by *Polq* shRNA + 10 μM M3814 (2.4% to 0.15%, *p* = 3.0 × 10^−4^) (Fig. 4d). Although the percentage of R−G+ cells was also decreased by *Polq* shRNA + 10 μM M3814 (7.8% to 2.8%, *p* = 0.0035), the R+G+ & R+G−/R−G+ ratio was significantly lower for *Polq* shRNA + 10 μM M3814 cells (0.31 vs 0.05, *p* = 2.6 × 10^−4^) (Fig. 4d, e). Additionally, the intensity of EGFP in the R−G+ population was significantly shifted from the weaker (monoallelic) to the stronger (biallelic) peak (Fig. 4f and Fig. S3). Taken together, BiPoD was able to tag endogenous genes.

**Figure 4.**
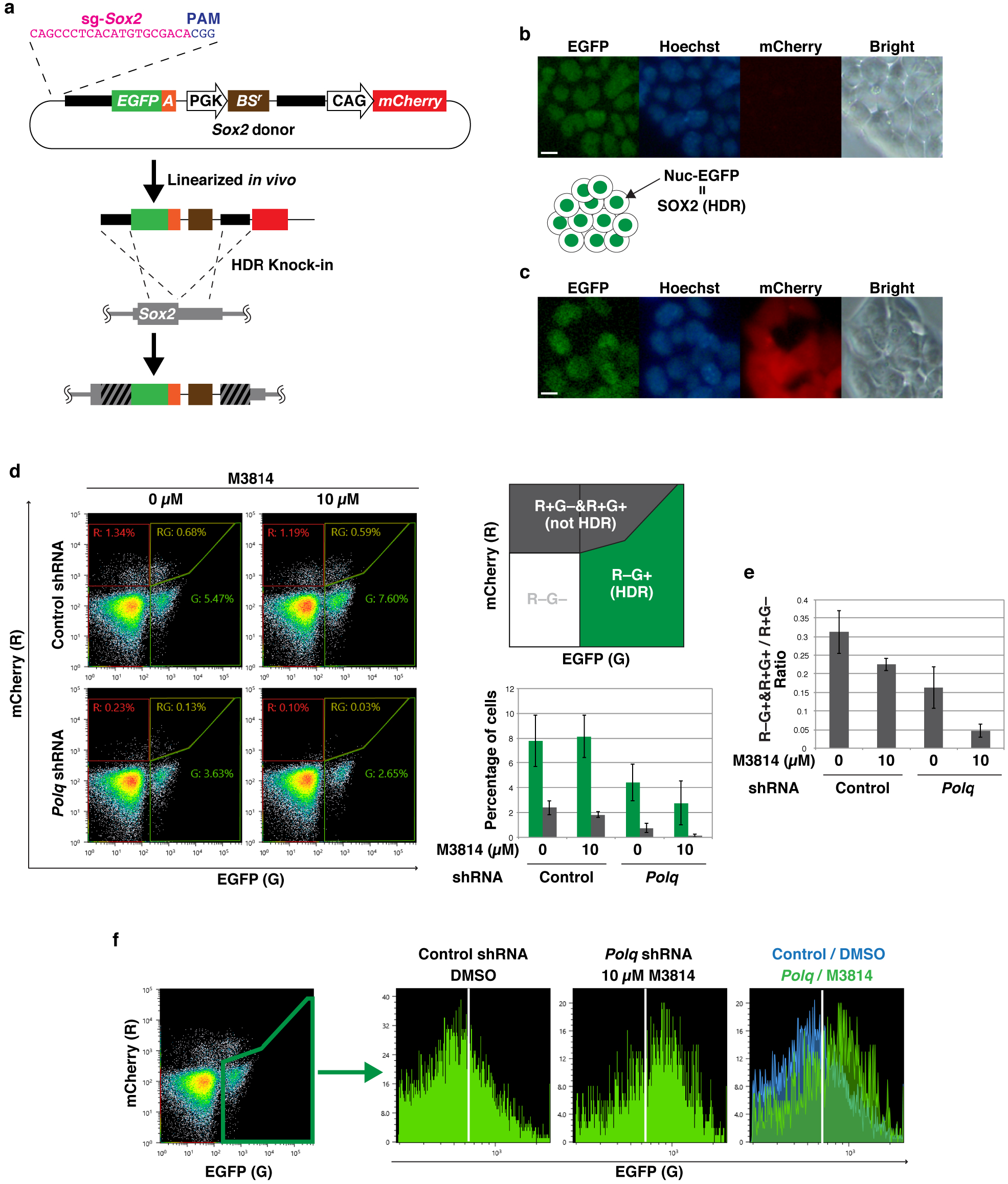
Tagging the endogenous *Sox2* gene with *Polq* knockdown and M3814 treatment. (a) Schematic illustration of the *Sox2* donor and the manner of HDR-mediated integration. Wide black bars and wide gray bars indicate the homology arms in the donor and the homologous region in the genome, respectively. Black-and-white striped bars denote the homologous region undergoing HDR. (b, c) Images of living cells expressing only EGFP (R−G+) (b) and both mCherry and EGFP (R+G+) (c) observed by fluorescence microscopy. Nuclei were stained with Hoechst 33342. Scale bar, 10 μm. (d) Flow cytometry analysis of mESCs transfected with the shRNA, *Sox2* donor, and Cas9/sgRNA expression vector. The cells were treated with DMSO or M3814 as illustrated in Fig. 2a. A series of experiments, from transfection to flow cytometric analysis, was independently performed in triplicate. Scatter plots show representative results from the three experiments. The average ± standard deviation percentages of R−G+ (green) and R+G+ & R−G+ (gray) cells are shown in the bar graph. (e) The ratio of R+G+ & R−G+ vs R−G+ is shown in the bar graph. (f) Fluorescence intensity of EGFP expressed by R−G+ cells. Representative histograms from three independent experiments are shown.

### Simultaneous biallelic knock-in into two different loci

Next, we attempted simultaneous knock-in of *Rosa26a-mCherry* and *Sox2-EGFP-mAID* (Fig. 5a). The donor DNA and Cas9/sgRNA expression vectors were the same as those used in the single knock-in experiments. *Polq* knockdown- or control-mESCs were simultaneously transfected with these four plasmids. To maintain the total amount of DNA transfected, the amounts of each DNA were half those used in the single knock-in experiments. *Polq* knockdown cells were treated with 10 μM M3814, while control cells (scrambled shRNA) were treated with DMSO. Transfected cells were treated with BS after Day 8. Surviving cells were predicted to harbor the *BS^r^* gene from the *Sox2* donor, regardless of the manner of integration. Isolated clones from surviving cells were expanded and analyzed by fluorescence microscopy and genomic PCR (Fig. 5b, c and Fig. S4). Only one control clone (scrambled shRNA + DMSO) (5%, 1/20) harbored knock-in alleles of *Rosa26a-mCherry* and *Sox2-EGFP-mAID*, both of which were monoallelic. Despite BS-resistance, 65% (13/20) of control clones did not harbor a knock-in *Sox2* allele. If the entire region of the *Sox2* donor is integrated in a non-HDR manner, mCherry is expressed as the negative selection marker. However, control clones 4 and 19 did not exhibit red fluorescence, probably because only part of the *Sox2* donor was integrated. In addition, PCR products of unexpected sizes were found in many control clones, indicating that large deletions or incorrect recombinations occurred in the target regions. In contrast, 96% (22/23) of *Polq* shRNA + 10 μM M3814 clones harbored a knock-in *Sox2* allele. Interestingly, 39% (9/23) of these clones harbored both *Rosa26a-mCherry* and *Sox2-EGFP-mAID*, and 26% (6/23) were biallelic at both loci. In contrast to the control, the fluorescence patterns of each *Polq* shRNA + 10 μM M3814 clone were almost completely consistent with those expected from the knock-in patterns (mCherry from *Rosa26a-mCherry* and Nuc-EGFP from *Sox2-EGFP-mAID*), and no unexpected PCR products were observed. These results demonstrated that BiPoD enabled efficient biallelic knock-in into two different loci simultaneously.

**Figure 5.**
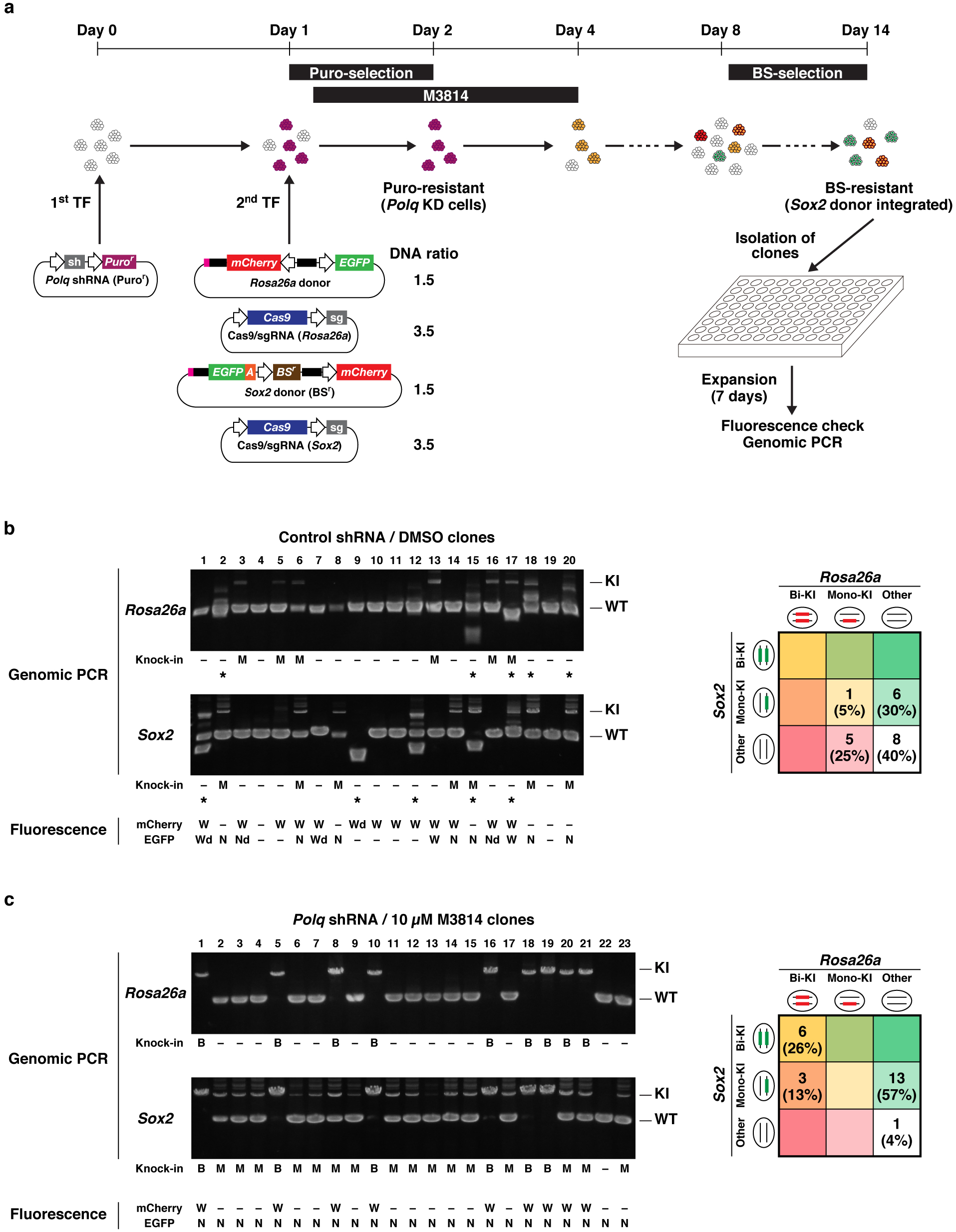
Simultaneous knock-in of *Rosa26a-mCherry* and *Sox2-EGFP-mAID*. Experimental scheme. Weight ratio of donor DNA and Cas9/sgRNA expression vector is shown. (b, c) Genomic PCR and fluorescence microscopy analysis of the clones. The PCR products were analyzed on agarose gels. Positions of knock-in alleles (KI) and wild-type alleles (WT) are indicated. Asterisks indicate the clones in which PCR bands of unexpected size were observed. Localization and intensity of mCherry and EGFP expressed in each clone are shown. W, whole (both cytoplasm and nucleus); N, nucleus; Wd, whole dim; Nd, nucleus dim; −, no fluorescence. The number and percentage of clones for each knock-in pattern are summarized in the table on the left.

### Treatment with novobiocin and M3814 increases biallelic knock-in efficiency

Novobiocin, a small-molecule antibiotic, was recently reported as an inhibitor of Pol θ [27]. We examined whether novobiocin can improve the efficiency of HDR-mediated knock-in in our scheme (Fig. 6a). When used in combination with 10 μM M3814, treatment with 300 μM novobiocin for 3 days completely killed mESCs, while a fraction of mESCs survived with 200 μM novobiocin. Flow cytometry analyses showed that 200 μM novobiocin + 10 μM M3814 decreased the proportion of R+G+ and R+G− cells, but the effect was not statistically different from that of 10 μM M3814 alone (Fig. 6b). These results indicated that novobiocin exerted a weaker effect compared with shRNA-mediated knockdown of *Polq*. However, 200 μM novobiocin + 10 μM M3814 increased biallelic knock-in, whereas there was almost no change for cells treated with novobiocin or M3814 alone (Fig. 6c and Fig. S5). Although the effect was weaker than that of shRNA-mediated knockdown of *Polq*, novobiocin exhibited synergistic effects with M3814, at least on the increase of biallelic knock-in.

**Figure 6.**
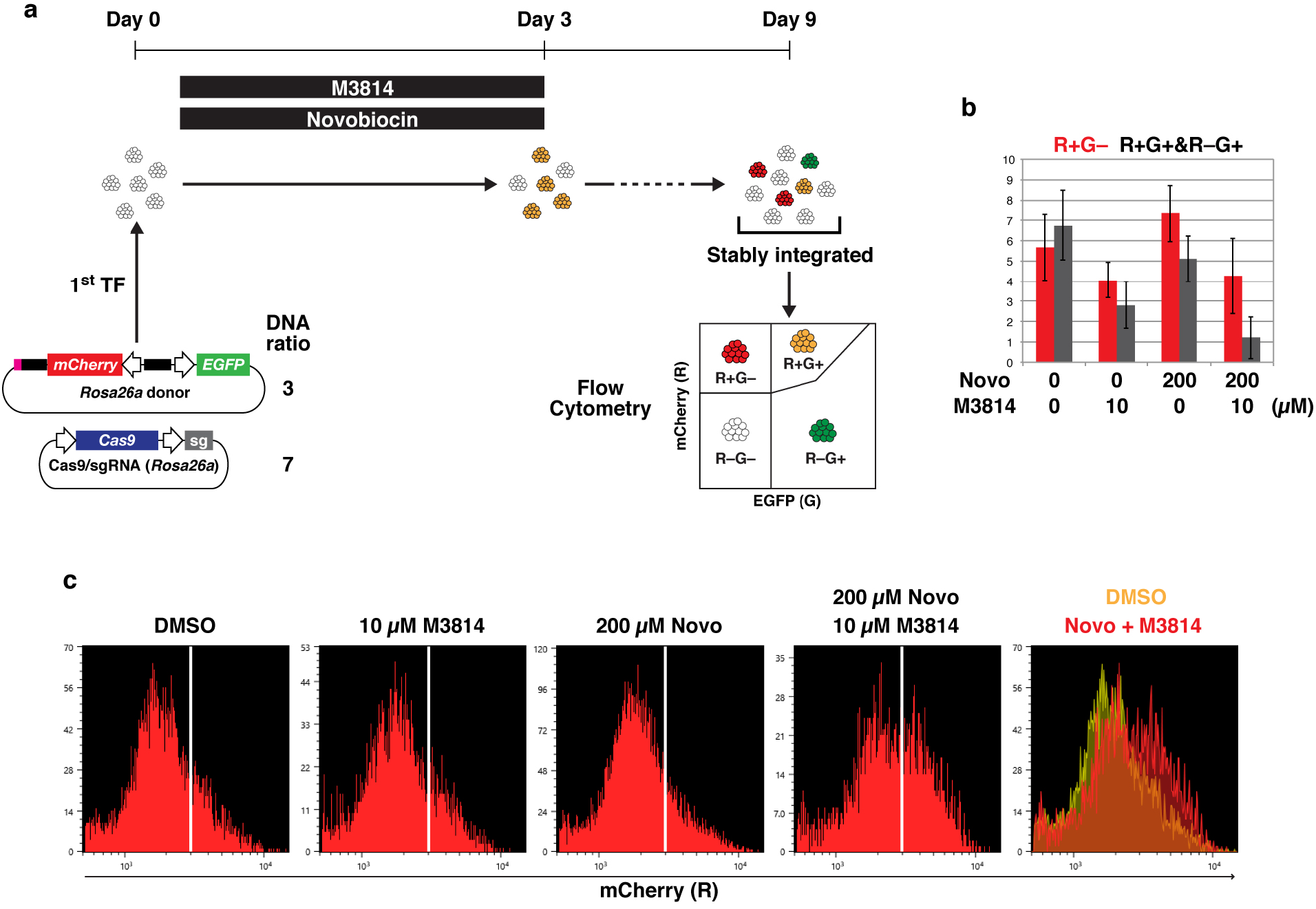
Effects of novobiocin and M3814 on HDR-mediated knock-in into *Rosa26a*. (a) Experimental scheme. Weight ratio of donor DNA and Cas9/sgRNA expression vector is shown. Flow cytometry analysis of mESCs transfected with the *Rosa26a* donor and Cas9/sgRNA expression vector. A series of experiments, from transfection to flow cytometric analysis, was independently performed in triplicate. The average ± standard deviation percentages of R+G− (red) and R+G+ & R−G+ (gray) cells are shown in the bar graph. (c) Fluorescence intensities of mCherry of R+G− populations in the experiments shown in Fig. 5c. Representative histograms from three independent experiments are shown.

## Discussion

Here, we propose BiPoD, an HDR-mediated knock-in method comprising *in vivo*-linearization of donors, knockdown of *Polq*, and inhibition of DNA-PK by M3814. Compared with the classic knock-in method, BiPoD is an improvement on two points: an increased rate of HDR by *in vivo*-linearization, and a decreased rate of non-HDR integration by inhibition of Pol θ and DNA-PK (Fig. 7). BiPoD is a simple and low-cost method: donor, expression of Cas9/sgRNA, and knockdown of *Polq* are all achieved by easily prepared plasmids. BiPoD enables easy and rapid establishment of biallelic knock-in mESCs.

**Figure 7.**
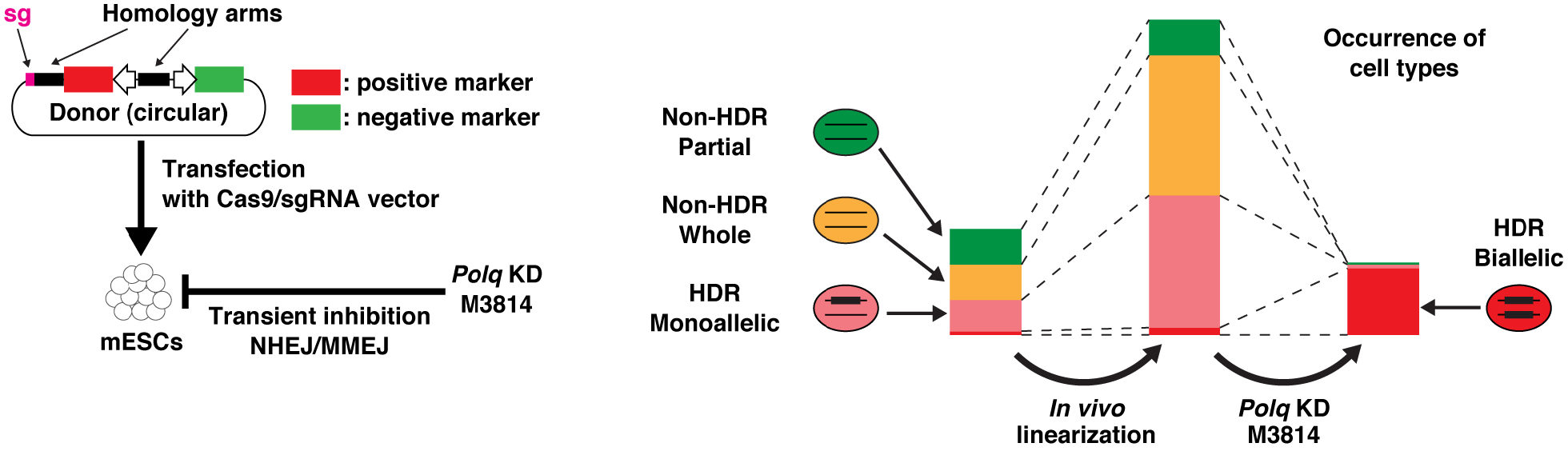
Illustration of BiPoD. The stacked bar graphs indicate the change in the occurrence of each cell type, based on the results of *Rosa26a-mCherry* knock-in.

Using the *Rosa26a* locus as a model, we show that *in vivo*-linearization of the donor increased the percentage of R+G− cells, mainly by HDR-mediated knock-in. Several studies have shown that *in vivo*-linearized donors are more efficiently integrated through HDR than non-linearized circular donors [28, 29]. In these studies, a CRISPR target site was located just outside the end of each homology arm. In contrast, we confirmed that a single CRISPR target site at the end of one homology arm was enough to significantly increase HDR-mediated knock-in, compared with donor linearization before transfection. An advantage of the single cut is that negative selection markers to detect unwanted integration via non-HDR pathways can be used. *In vivo*-linearization also increased the percentage of R+G+ cells, probably generated by integration of all or almost all regions of the donor. The design of the *Rosa26a* donor, in which the single CRISPR target sequence was located outside one of the homology arms, is similar to that used in a previous report [9]. They showed that the entire donor DNA sequence is efficiently integrated into the Cas9-induced DSB site when the donor had one CRISPR target sequence. Similarly, on-target integration of the entire donor DNA sequence probably increased the R+G+ population in our study. However, the R−G+ population was not increased by *in vivo*-linearization, indicating that the occurrence of random integrations was not substantially affected.

Recent studies have shown that simultaneous inhibition of Pol θ and NHEJ-related factors abolishes random integration through NHEJ and MMEJ [12, 13]. These studies used constitutive knock-out of genes encoding Pol θ and subunits of LIG4. We demonstrated that transient knockdown of *Polq* by shRNA was also effective, making the strategy applicable for general knock-in experiments. Inhibition of Pol θ was important, particularly for suppressing partial integration of donor DNA. Another important advance concerns DNA-PK inhibitors. NU7441 failed to completely abolish non-HDR integrations in *Polq* knock-out or *Polq* knockdown mESCs (Ref. 12 and Fig. 2c); however, this was achieved using M3814. The effect of M3814 alone on knock-in into both *Rosa26a* and *Sox2* loci was not significant; however, the combination of M3814 and *Polq* knockdown almost completely abolished non-HDR integration events, without preventing HDR. Accordingly, large deletion and incorrect integration at the target loci, probably mediated by NHEJ and MMEJ, were completely suppressed by M3814 + *Polq* knockdown (Fig. 3b and Fig. 5b, c).

M3814 + *Polq* knockdown dramatically improved the efficiency of biallelic knock-in. In this study, 96% of R+G− cells (i.e. 4.2% of all cells) harbored biallelic *mCherry* genes in *Rosa26a* (Fig. 2d and 3c). This high efficiency of BiPoD enables biallelic knock-in cells to be obtained without the need for marker genes during clone isolation. Furthermore, BiPoD achieved simultaneous biallelic knock-in at two different loci. In this study, only one marker gene (*BS^r^*) was used for selection of stably integrated cells. Consequently, 26% of selected cells were double-biallelic. Thus, double-biallelic knock-in mESCs can be established easily and quickly by BiPoD using only one selectable marker. The increased biallelic knock-in efficiency is likely to result from inhibition of both NHEJ and MMEJ pathways. Generally, NHEJ/MMEJ repairs CRISPR/Cas9-induced DSB sites faster than HDR and causes indel mutations or insertion of donor. Such sites are no longer targeted by CRISPR/Cas9 and subsequent HDR. However, DSB sites have to be repaired by HDR using foreign donors or undamaged chromosomes as templates when both NHEJ/MMEJ are inactivated, although sites repaired using undamaged chromosomes recover their original genomic sequence and can, therefore, be cleaved by CRISPR/Cas9 again. Consequently, all target alleles are eventually cleaved and repaired by HDR using donor DNAs, resulting in biallelic knock-in or apoptosis.

During the knock-in experiments, several types of partial integration were observed. Knock-in into *Rosa26a* generated a minor population of cells that only expressed a negative selection marker (R− G+) (Fig. 2). Simultaneous knock-in into *Rosa26a* and *Sox2* generated some clones that harbored randomly integrated *Sox2* donors, but that did not express the negative expression marker (Fig. 5b). Such partial integrations create false-positives and prevent rapid acquisition of desired cell lines. These results also indicate that a similar amount of silent integration, which is undetectable by marker expression, occurs concomitantly and silently disrupts the genome. Knock-in into *Rosa26a* showed that *in vivo*-linearization did not increase the percentage of R−G+ cells, which decreased the chance of obtaining cells harboring partial integrations. Furthermore, functional inhibition of Pol θ significantly decreased partial integration, and *Polq* knockdown combined with M3814 treatment abolished almost all partial integrations. Therefore, BiPoD facilitates and accelerates successful acquisition of scarless knock-in cells by excluding partial integrations.

BiPoD requires a two-step transfection protocol, followed by antibiotic-selection of Pol θ-downregulated cells, limiting the application of this method. Chemical inhibitors against Pol θ are a promising solution to this problem. Novobiocin is the only published small compound that inhibits Pol θ [27]. We found that novobiocin + M3814 improved biallelic knock-in efficiency compared with M3814 alone. However, the effects of novobiocin were clearly weaker than those of shRNA knockdown, even if used at the maximum sub-toxic concentration. This is likely to be because of the specificity of the activity of novobiocin. Novobiocin was used at 50-200 μM to suppress Pol θ activity in U2OS cells [27]; however, novobiocin weakly inhibits eukaryotic topoisomerase II and HSP90 [30, 31]. Partial inhibition of these extra target proteins might potentiate the cytotoxicity of novobiocin, preventing sufficient inactivation of Pol θ. Pol θ is a promising target for cancer therapy; therefore, Pol θ inhibitors are currently in development in multiple companies and laboratories [32]. More specific and potent Pol θ inhibitors will enable a one-step transfection BiPoD protocol that does not need antibiotic selection.

## Materials and Methods

### Reagents

NU7441 and M3814 were purchased from Cayman Chemical (Ann Arbor, MI). These reagents were dissolved in DMSO and used with a final DMSO concentration of 0.1%. Puromycin (FUJIFILM Wako Pure Chemical Corporation, Osaka, Japan) and Blasticidin (MP Biomedicals, Santa Ana, CA) were dissolved in sterilized water and used for selection at 1 μg/mL and 10 μg/mL, respectively. Novobiocin sodium (Kanto Chemical Co., Inc., Tokyo, Japan) was dissolved in sterilized water and used at the indicated concentrations.

### Plasmids

pX330-U6-Chimeric_BB-CBh-hSpCas9 (a gift from Feng Zhang, Addgene #42230) was used for expression of Cas9/sgRNA. The CRISPR target sequences were as follows: *Rosa26a*, 5′-ACTCCAGTCTTTCTAGAAGA-3′ [22] and *Sox2*, 5′-CAGCCCTCACATGTGCGACA-3′ [33]. The *Rosa26a* donor vector was based on pROSA26-PA (a gift from Frank Costantini, Addgene #21271). The *mCherry* expression cassette from pCAGGS-mCherry (a gift from Phil Sharp, Addgene #41583) was inserted between the homology arms. The guide RNA sequence targeting *Rosa26a* with a PAM sequence was inserted into the *Kpn*I and *Hind*III sites, while the EGFP expression cassette was inserted into the *Sac*I and *Nco*I sites. The *Sox2* donor vector was constructed as follows: *EGFP-mAID* and the PGK-*BSD* expression cassette were flanked by the homology arms for *Sox2*. The *mAID*-PGK-*BSD* sequence was derived from pMK391 (a gift from Masato Kanemaki, RIKEN BRC #RDB16819 [34]). Silent mutations were introduced into the 5′ homology arm to prevent targeting by CRISPR/Cas9. A stretch of sequence was inserted upstream of the pCAGGS-mCherry CAG promoter. Detailed maps of the donor plasmids are shown in Fig. S6 and S7. The puromycin-selectable shRNA expression vector (a gift from Jun Ohgane) is the same as that described previously [35]: a pENTR-based shRNA expression vector (a gift from Makoto Miyagishi and Hideo Akashi) with a puromycin resistance gene expression cassette followed by an internal ribosome entry site (IRES) and *Venus* gene [36].

### Cell culture

The J1 mESC line was purchased from American Type Culture Collection. mESCs were cultured on 0.1% gelatin-coated dishes in KnockOut DMEM (Thermo Fisher Scientific, Waltham, MA) supplemented with 15% fetal bovine serum (Biowest, Nuaillé, France), 1000 U/mL leukemia inhibitory factor (Merck Millipore, Burlington, MA), 1% L-glutamine (FUJIFILM Wako Pure Chemical Corporation), 1% non-essential amino acids (FUJIFILM Wako Pure Chemical Corporation), 0.18% 2-mercaptoethanol (Thermo Fisher Scientific), and 1% penicillin-streptomycin (FUJIFILM Wako Pure Chemical Corporation). Leukemia inhibitory factor was added at 2000 U/mL during DNA-PK inhibitor treatment and simultaneous knock-in experiments.

### Transfection

Transfection was performed using Lipofectamine 2000 (Thermo Fisher Scientific) and the single suspension method [37]. Briefly, trypsinized mESCs were collected in a 1.5 mL tube. After removal of supernatant, cells were resuspended in DNA-Lipofectamine 2000 complex in Opti-MEM (Thermo Fisher Scientific) by pipetting, then incubated at room temperature for 5 min. The mESCs were then diluted in mESC medium (without penicillin-streptomycin) and transferred to a gelatin-coated plate for culture. When the transfectants were treated with DNA-PK and/or Pol θ inhibitors, the medium was replaced with fresh medium containing the inhibitors 4 hours after transfection.

### Flow cytometry

Fluorescence of living cells was analyzed using an SH800 Cell Sorter (Sony, Tokyo, Japan).

### Knockdown of *Polq*

The shRNA target sequences were 5′-GAATATGAGTGATAGTATACT-3′ (*Polq*) and 5′-CAACAAGATGAAGAGCACCAA-3′ (scrambled). To confirm knockdown, the puromycin-selected transfectants were directly subjected to RT-qPCR analysis using the CellAmp™ Direct TB Green^®^ RT-qPCR Kit (TaKaRa Bio Inc., Shiga, Japan) at 3 days after transfection. The primers for RT-qPCR were as follows: *Polq*, 5′-GACTCTGGGTTCCACCAGAAG-3′ and 5′-GGCCTGGTAAAGGATGCAAG-3′, *Actb*, 5′-TTCTACAATGAGCTGCGTGTGG-3′ and 5′-ATGGCTGGGGTGTTGAAGGT-3′.

### Genotyping of *Rosa26a* knock-in clones

R+G− cells were collected by fluorescence activated cell sorting (FACS) using an SH800 Cell Sorter (Sony) and seeded on gelatin-coated dishes at low density to obtain clonal colonies. After culturing for 4 days, colonies were picked and transferred to a 96-well plate. During picking, we ensured that isolated colonies were not contaminated with other cells. On the following day, the cells were observed by fluorescence microscopy to confirm whether they were R+G−. R−G− clones found here were excluded from subsequent experiments. We also confirmed that there was no contamination of R−G−, R−G+, or R+G+ cells in the R+G− clones. The R+G− clones were expanded for 6-7 days. Their fluorescence was observed again just before genomic DNA extraction to confirm that all cells were R+G−. Then, cells were lysed in 0.1 M Tris-HCl buffer (pH 8.0) containing 0.2% sodium lauryl sulfate, 5 mM ethylenediaminetetraacetic acid-2Na, 200 mM NaCl, and 5 μg/mL Proteinase K (FUJIFILM Wako Pure Chemical Corporation). Genomic DNA for each clone was extracted by isopropanol precipitation followed by rinsing with 70% ethanol and dissolving in 0.1× Tris-EDTA buffer (Thermo Fisher Scientific). PCR was performed with KOD FX Neo (TOYOBO, Osaka, Japan) according to the manufacturer’s instructions. The primers for amplification of the *Rosa26a* locus were 5′-CTCCGGCTCCTCAGAGAGCCTCGGCTAGGTAG-3′ (outside the 5′ arm) and 5′-GGAAAATACTCCGAGGCGGATCACAAGCAATAATAACCTGTAG-3′ (inside the 3′ arm).

### Genotyping of double knock-in clones

Cloning and analysis were performed as for *Rosa26a* knock-in clones with several modifications. The transfectants were treated with 10 μg/mL BS for 2 days, then passaged at low density. After culturing for 4 days with 10 μg/mL BS, colonies were picked into a 96-well plate, cultured for 5 days, passaged into a 48-well plate and further cultured for 2 days. Genomic DNA was then extracted. Fluorescence was observed at 3 days after colony picking and just before genomic DNA extraction to identify and exclude contaminated clones. The primers for amplification of the *Sox2* locus were 5′-GAGAACCCCAAGATGCACAACTCGGAGATCAGCAAG-3′ (outside the 5′ arm) and 5′-CATGGATTCTCGGCAGCCTGATTCCAATAACAGAG-3′ (outside the 3′ arm).

### Microscopy

Living mESCs were observed with a fluorescence IX71 microscope (Olympus, Tokyo, Japan) at 20× magnification. Nuclei were stained with Hoechst 33342 (Dojindo Laboratories, Kumamoto, Japan).

### Statistics

All experiments were repeated three times. Statistical significance was determined by Student’s *t*-test (two samples) or Tukey’s multiple comparison test (all pairs from ≥ 3 samples) using R software. All P-values from Tukey’s multiple comparison tests are shown in Supplementary Tables 1-4.

## Acknowledgements

We thank Dr. Jun Ohgane for providing the shRNA expression vector, Dr. Tetsuya S. Tanaka for providing the IRES-Venus containing vector, Drs. Makoto Miyagishi and Hideo Akashi for providing the pENTR vector, Dr. Feng Zhang for providing the pX330 vector, Dr. Frank Costantini for providing the pROSA26-PA vector, Dr. Phil Sharp for providing pCAGGS-mCherry, and Dr. Masato Kanemaki for providing the pMK391 vector. We thank Jeremy Allen, PhD, from Edanz Group (https://en-author-services.edanz.com/ac) for editing a draft of this manuscript. This work was supported by JSPS Grants-in-Aid for Scientific Research, Grant Numbers 20K06449, 18H03993, 18H02100, 26221204, and a Waseda University Grant for Special Research Projects (2020C-289).

## Author Contributions

D.A. and Y.N. conceived the idea. D.A. performed all the experiments. D.A. and Y.N. wrote the manuscript.

## Data Availability

The datasets generated during and/or analyzed during the current study are available from the corresponding author on reasonable request.

