## Supplementary Figures for "Efficient biallelic knock-in in mouse embryonic stem cells by *in vivo*-linearization of donor and transient inhibition of DNA Polymerase θ/DNA-PK"

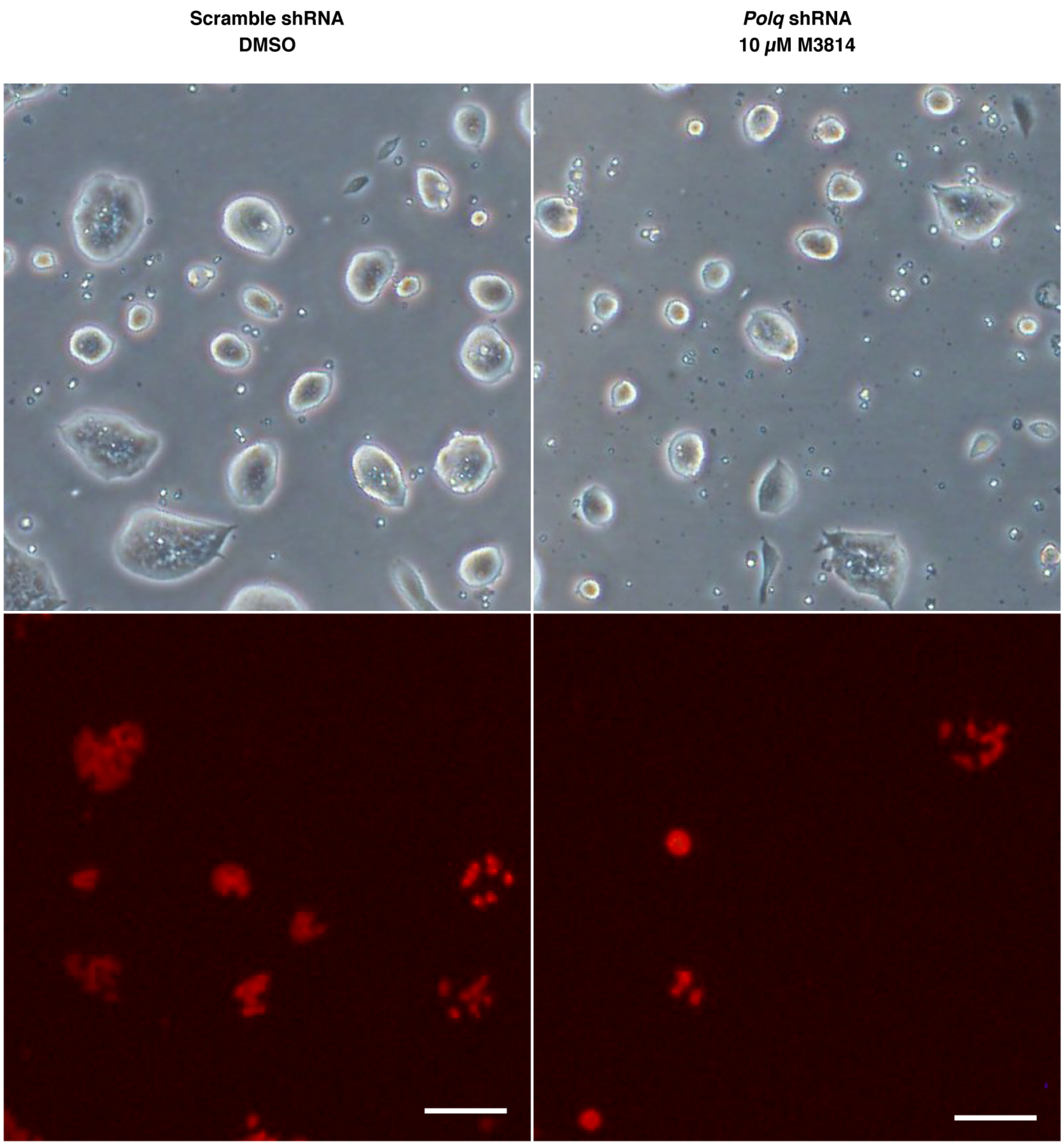


Supplementary figure 1.

Images of control and *Polq* shRNA + 10 μM M3814 cells just before flow cytometry analysis related to Fig. 2d. Scale bar, 100 μm.

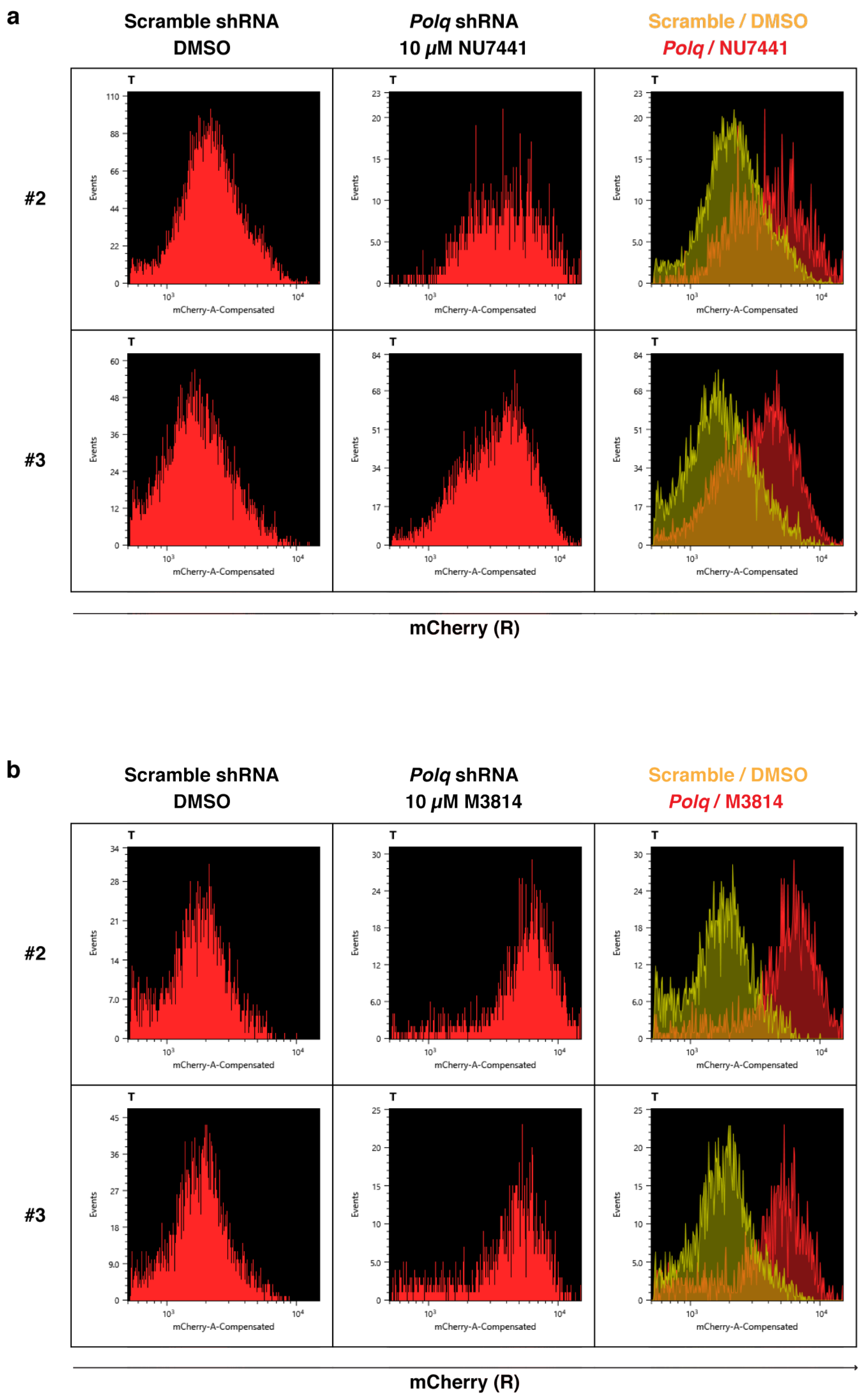


Supplementary figure 2.

Fluorescence intensities of mCherry in R+G– populations from *Polq* shRNA + 10 µM NU7441 (a) and *Polq* shRNA + 10 µM M3814 (b) cells (results not shown in Fig. 3a).

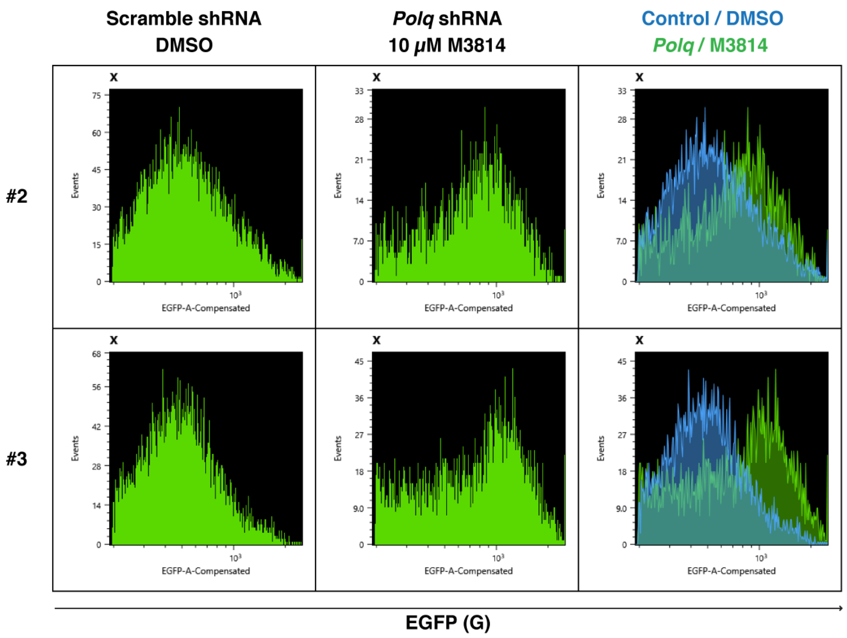


Supplementary figure 3.

Fluorescence intensities of EGFP in R–G+ populations from *Polq* shRNA + 10 μM M3814 cells (results not shown in Fig. 4f).

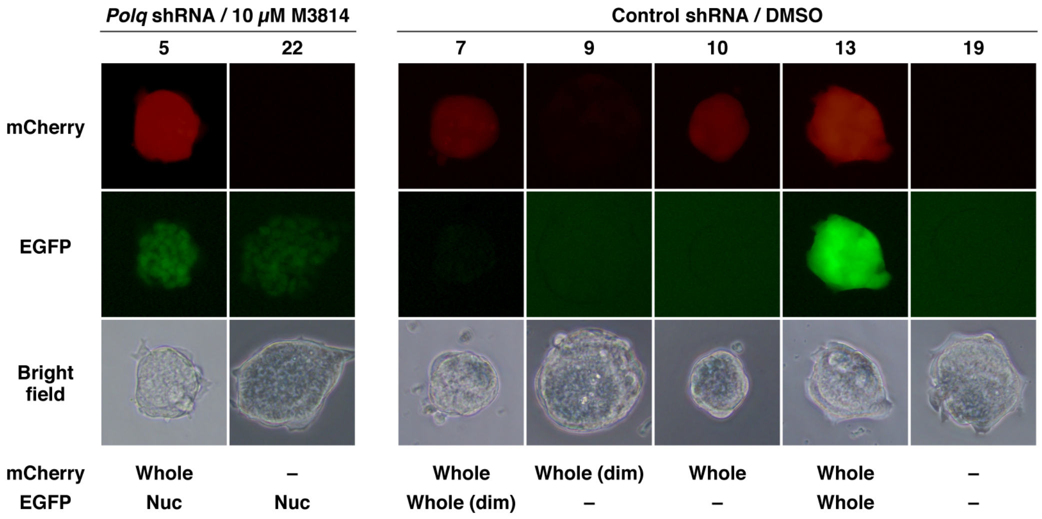


Supplementary figure 4.

Images of isolated clones observed by fluorescence microscopy related to Fig. 5.

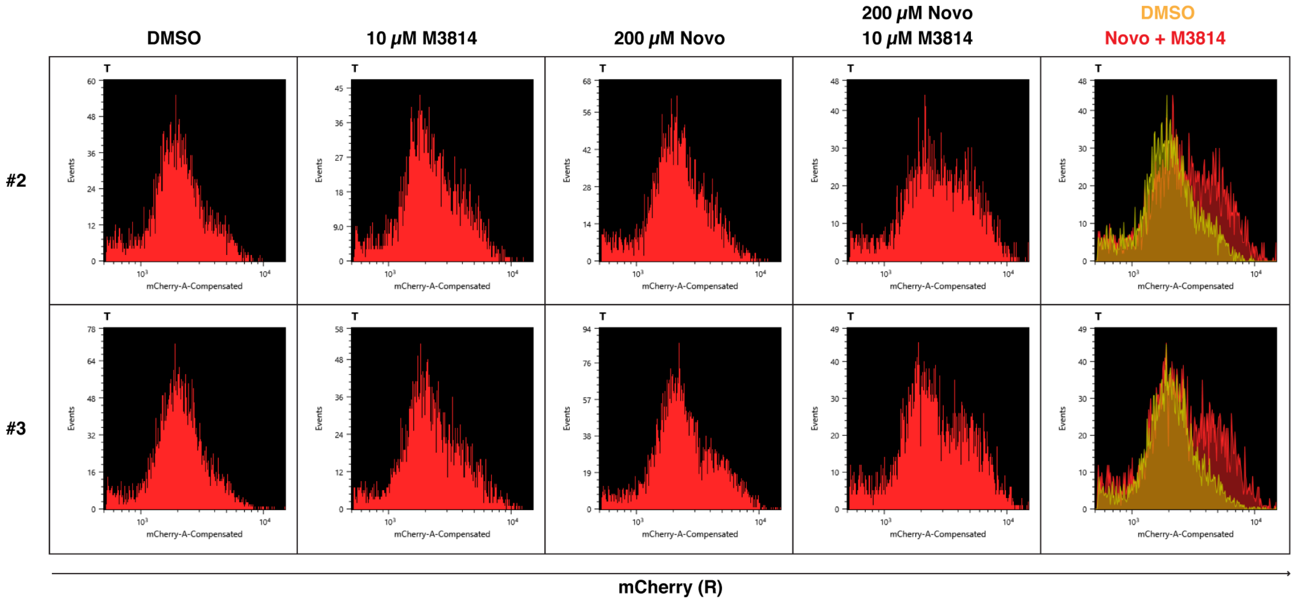


Supplementary figure 5.

Fluorescence intensities of mCherry in R+G– populations from the experiments shown in Fig. 6 (results not shown in Fig. 6c).

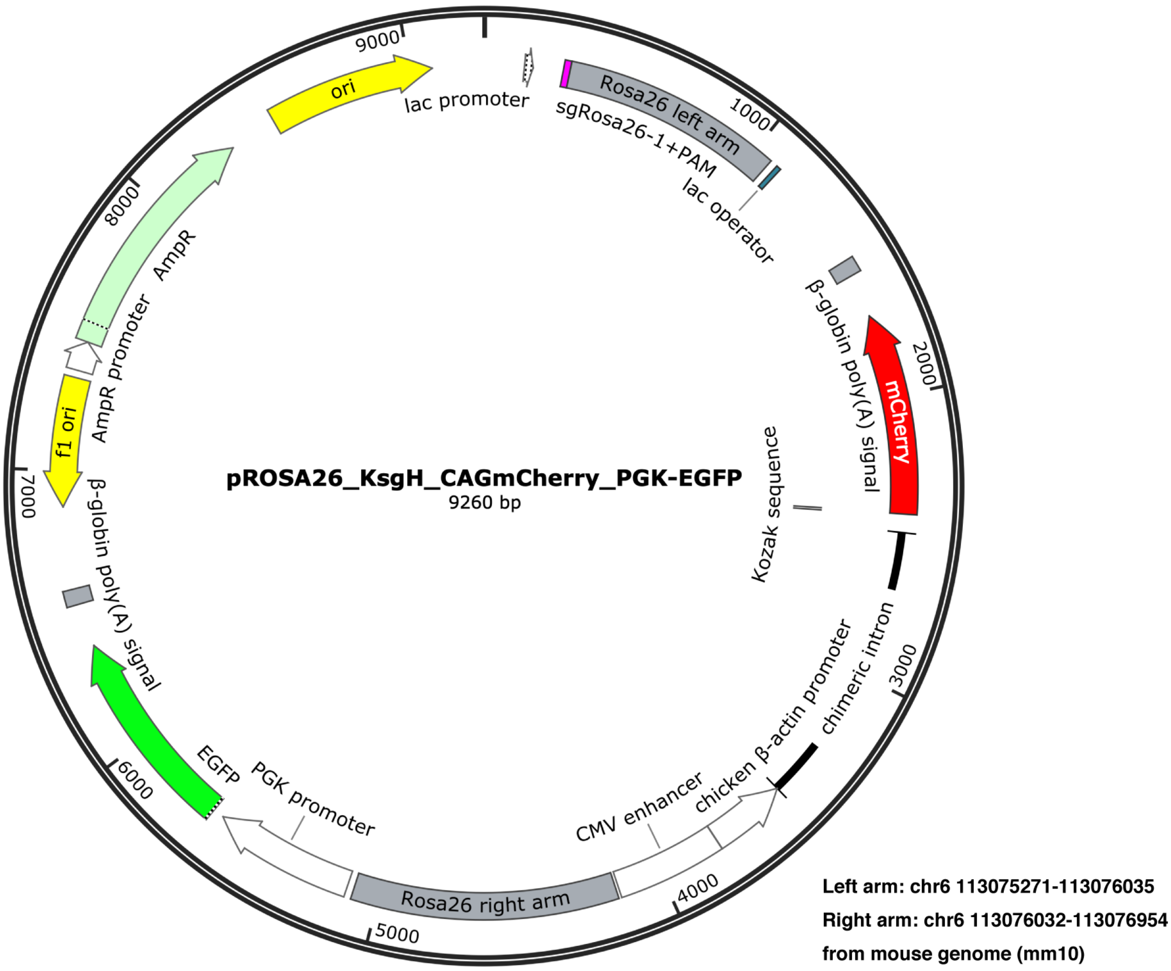


Supplementary figure 6.

Map of the donor plasmid for *Rosa26a-mCherry*.

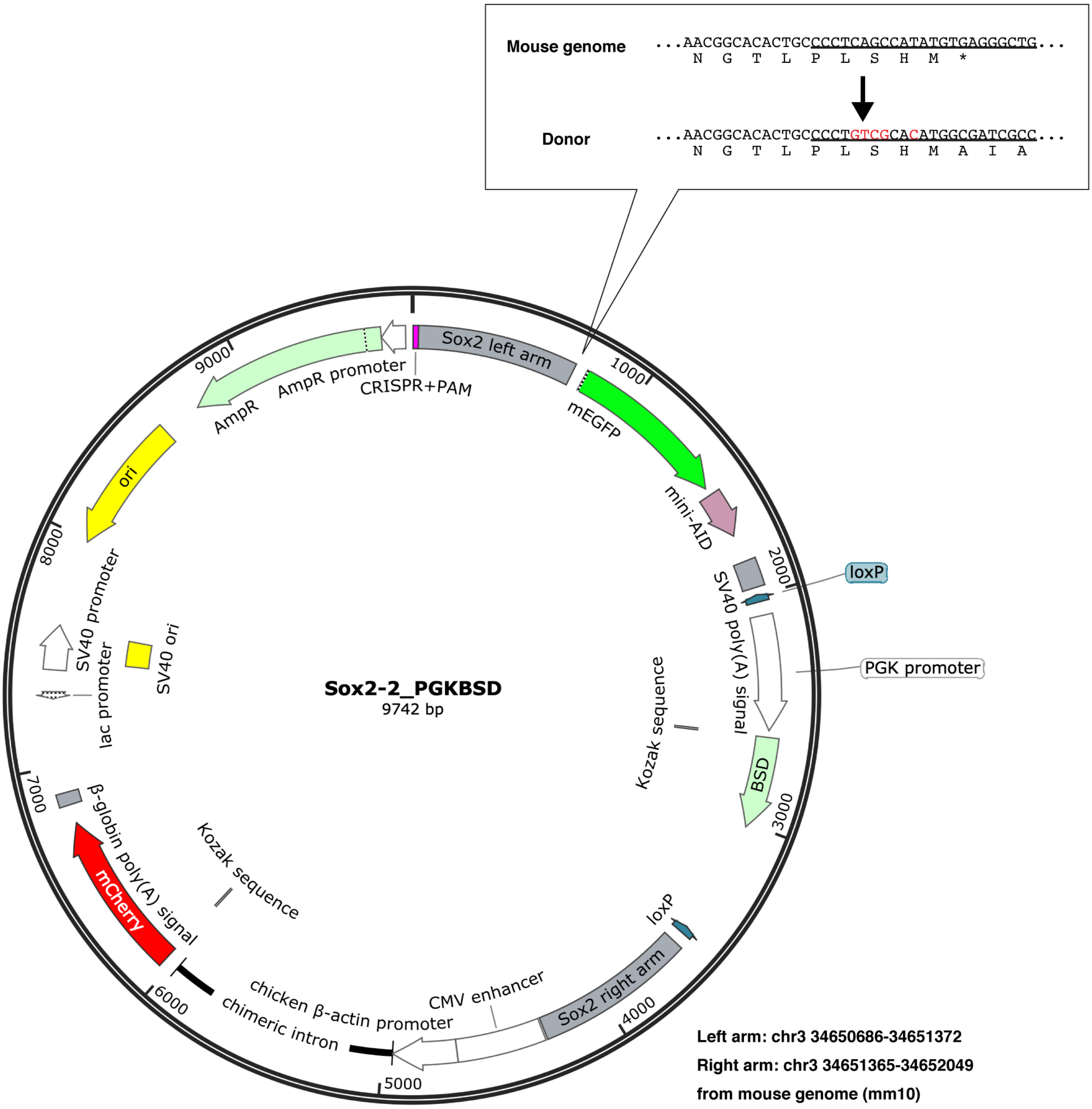


Supplementary figure 7.

Map of the donor plasmid for *Sox2-EGFP-mAID*. The sequence around the *Sox2* stop codon is shown. The target sequence for CRISPR/Cas9 in the mouse genome is underlined.

**Supplementary information. Uncropped images with size marker indications, related to Fig.3c.**

Control shRNA
DMSO

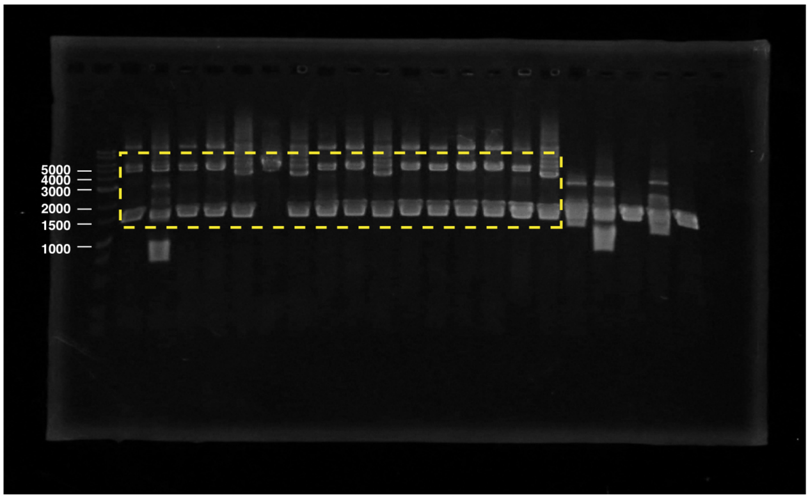


*Polq* shRNA
10 $\mu$M NU7441

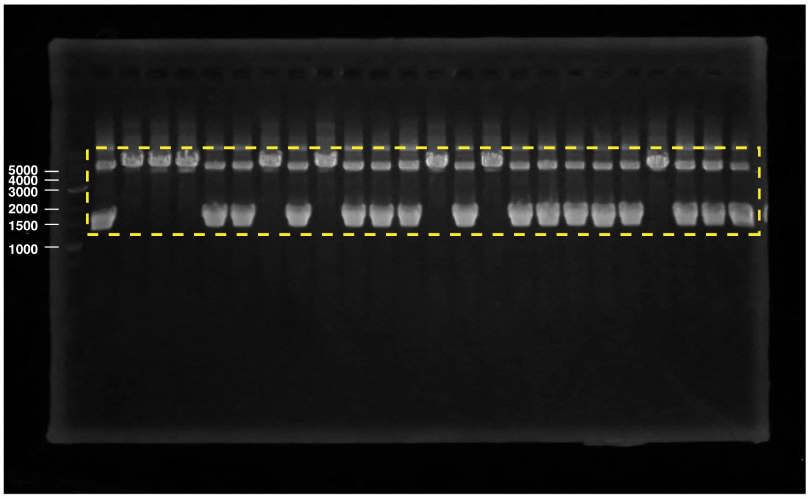


*Polq* shRNA
10 $\mu$M M3814

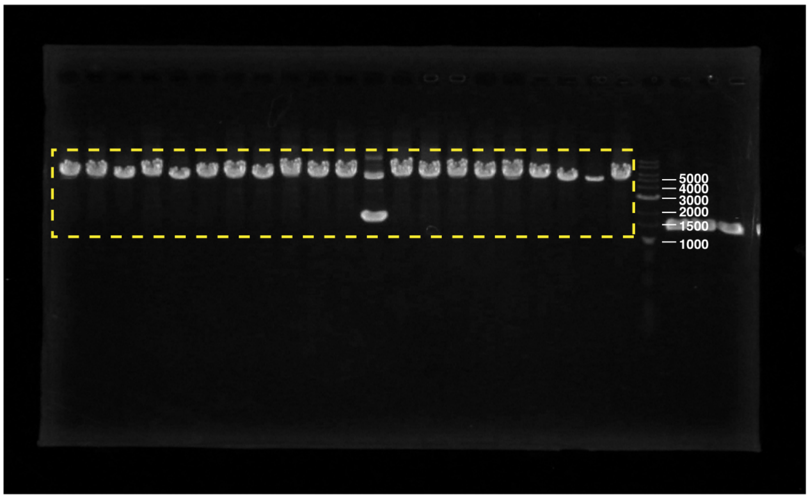

Supplementary information. Uncropped images with size marker indications, related to Fig.5b and c.

*Rosa26a*

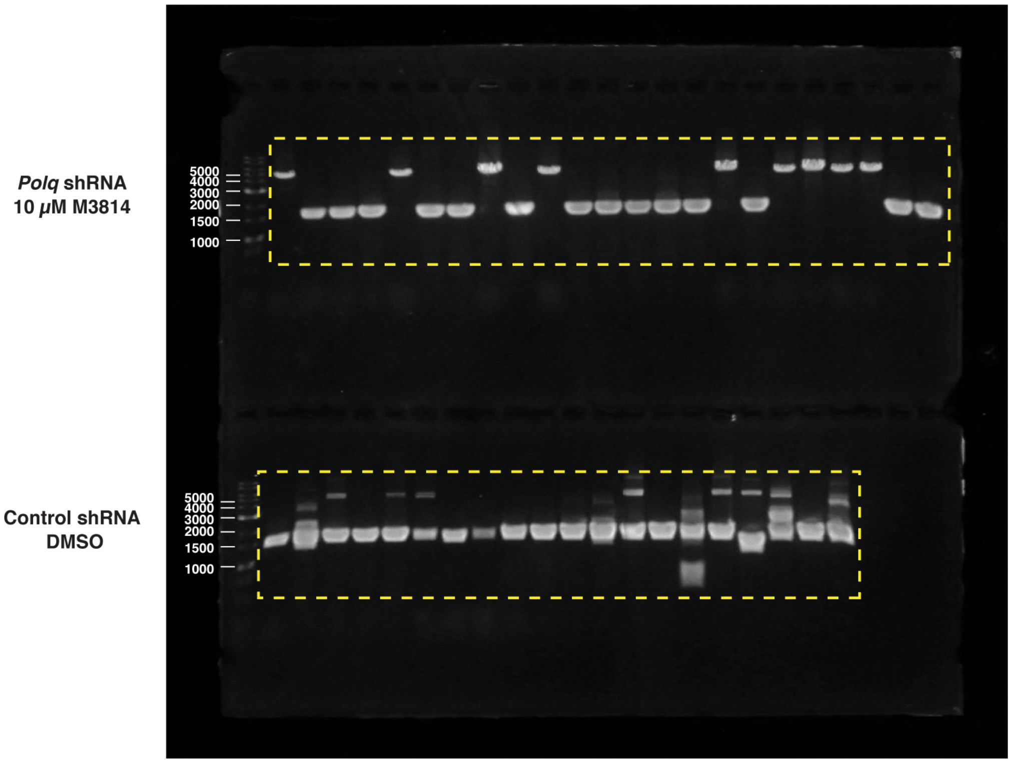


*Sox2*

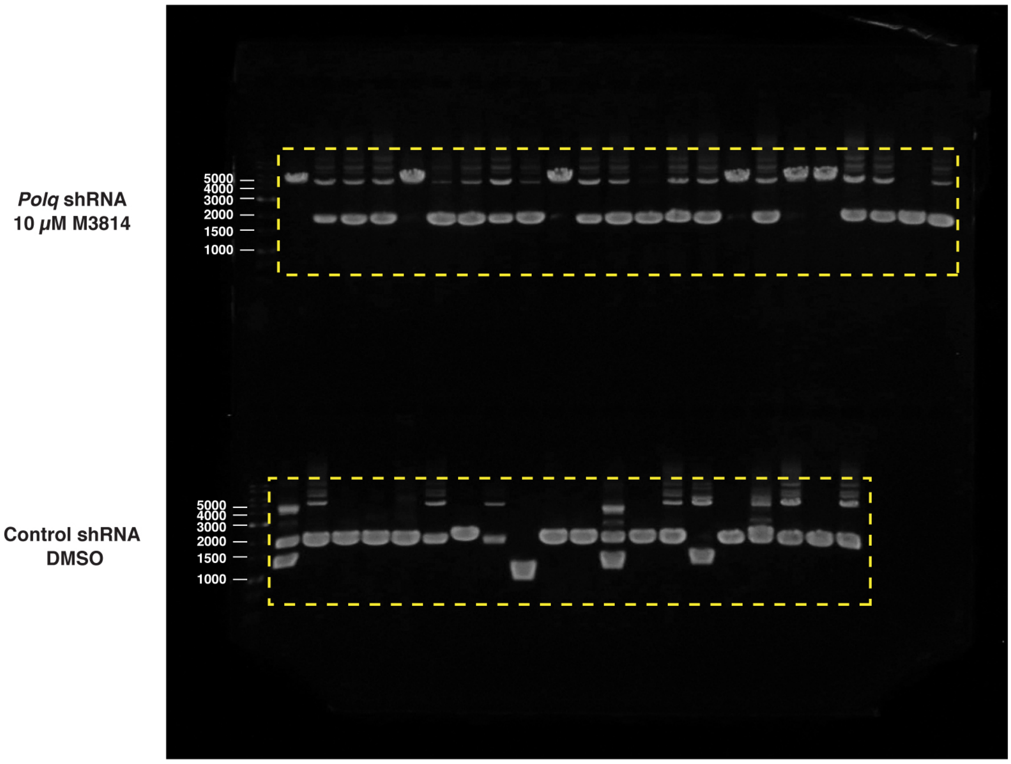
